# Mild SARS-CoV-2 infection in rhesus macaques is associated with viral control prior to antigen-specific T cell responses in tissues

**DOI:** 10.1101/2022.01.06.475282

**Authors:** Christine E. Nelson, Sivaranjani Namasivayam, Taylor W. Foreman, Keith D. Kauffman, Shunsuke Sakai, Danielle E. Dorosky, Nickiana E. Lora, NIAID/DIR Tuberculosis Imaging Program, Kelsie Brooks, E. Lake Potter, Mario Roederer, Alan Sher, Daniela Weiskopf, Alessandro Sette, Emmie de Wit, Heather D. Hickman, Jason M. Brenchley, Laura E. Via, Daniel L. Barber

**Author notes:** Corresponding author: Daniel L. Barber. Members of the NIAID/DIR Tuberculosis Imaging Program include: Ayan Abdi, Emmuanual K. Dayao, Joel D. Fleegle, Felipe Gomez, Michaela K. Piazza, Katelyn M. Repoli, Becky Y. Sloan, Ashley L. Butler, April M. Walker, Danielle M. Weiner, Michael J. Woodcock, and Alexandra Vatthauer.

## Abstract

SARS-CoV-2 primarily replicates in mucosal sites, and more information is needed about immune responses in infected tissues. We used rhesus macaques to model protective primary immune responses in tissues during mild COVID-19. Viral RNA levels were highest on days 1-2 post-infection and fell precipitously thereafter. ^18^F-fluorodeoxyglucose (FDG)-avid lung abnormalities and interferon (IFN)-activated myeloid cells in the bronchoalveolar lavage (BAL) were found on days ∼3-4. Virus-specific effector CD8 and CD4 T cells were detectable in the BAL and lung tissue on days ∼7-10, after viral RNA, lung inflammation, and IFN-activated myeloid cells had declined. Notably, SARS-CoV-2-specific T cells were not detectable in the nasal turbinates, salivary glands, and tonsils on day 10 post-infection. Thus, SARS-CoV-2 replication wanes in the lungs prior to T cell responses, and in the nasal and oral mucosa despite the apparent lack of Ag-specific T cells, suggesting that innate immunity efficiently restricts viral replication during mild COVID-19.

**ONE SENTENCE SUMMARY:** SARS-CoV-2 infection leads to mild, focal lung inflammation, and type I IFN activated myeloid cells that mostly resolve prior to the influx of virus-specific effector T cells or antibody responses in rhesus macaques.

## INTRODUCTION

SARS-CoV-2 infection has a spectrum of clinical outcomes, ranging from asymptomatic to fatal. There is a need to parse out the role of individual immune cell types and molecular pathways that contribute to effective control of viral infection in asymptomatic/mild disease and those leading to organ failure during severe COVID-19. Increased pro-inflammatory cytokines^1–4^, deficient type I interferon (IFN) responses^5–7^, activation of inflammasomes^8^, neutrophils^9–11^, and monocytes/macrophages^1, 2, 11–14^ have all been associated with severe COVID-19. Coordinated activation of CD8 and CD4 T cells, T cell activation state, and antigen (Ag)-specificity have all been linked to favorable outcomes of SARS-CoV-2 infection^15–20^. While neutralizing antibodies are clearly protective in immune hosts, T cell responses may also contribute to the protection provided by vaccination and natural infection^21–24^.

Most studies of immune correlates of COVID-19 disease severity in humans have focused on sampling of peripheral blood. Nevertheless, some studies have observed infiltration of immune cells into the BAL fluid, post-mortem lung tissue acquired from fatal COVID-19 cases, or from individuals undergoing medically necessary procedures^12, 14, 25, 26^. Notably, on autopsy several reports have observed a surprising lack of immune cells infiltrating into extrapulmonary tissues despite the prescience of high levels of virus^27–31^. These data highlight the importance of understanding the early host response in pulmonary and extrapulmonary tissues in the first few days after SARS-CoV-2 infection.

Animal models can be employed to obtain a detailed understanding of the host response in infected tissues. Studies in SARS-CoV-2 susceptible species provide insights into COVID-19 disease pathogenesis. For example, transgenic mouse strains expressing human angiotensin converting enzyme (ACE)2 (K18-hACE2), and mice induced to express hACE2 with viral vectors, are highly susceptible to SARS-CoV-2 infection^32–36^. Syrian hamsters and ferrets are also moderately susceptible and shed infectious virus^37–41^. On the other hand, species more resistant to SARS-CoV-2 disease are useful tools in examining mechanisms of efficient control of viral replication. Several non-human primate (NHP) species can be experimentally infected with SARS-CoV-2^42, 43^. Rhesus macaques, cynomolgus macaques, and African green monkeys typically develop mild signs after SARS-CoV-2 infection^44–54^. SARS-CoV-2 immune and vaccinated rhesus macaques are protected from reinfection primarily by neutralizing antibodies and to a lesser extent anamnestic T cell responses^55–65^. Thus, NHP are suitable for the study of protective host immune responses associated with mild SARS-CoV-2 infection.

In this study, we use the rhesus macaque model of mild COVID-19 to examine the (1) kinetics of lung inflammation using ^18^FDG positron emission tomography computed tomography (PET/CT) imaging, (2) innate immune responses using single cell RNA sequencing (scRNAseq), and (3) the tissue distribution of SARS-CoV-2-specific T cell responses by flow cytometry. Our findings suggest that mild SARS-CoV-2 disease and efficient control of the infection are temporally correlated with activation of myeloid cells by type I IFN, prior to the induction of Ag-specific T and B cell responses. Moreover, they reveal a strong propensity for Ag-specific T cell migration into the pulmonary compartment compared to other mucosal sites of infection.

## RESULTS

### Radiologic and virological outcomes of SARS-CoV-2 infection in rhesus macaques

Six, male rhesus macaques were infected with SARS-CoV-2/USA-WA-1 at 1x10^6^ tissue culture infectious dose (TCID)_50_ intranasally (i.n.) and 1x10^6^ TCID_50_ intratracheally (i.t.), for a total dose of 2x10^6^ TCID_50_ (Table 1). ^18^FDG PET/CT imaging showed evidence of heterogeneous inflammatory foci with increased ^18^FDG uptake (Fig 1A, B) and lesion density (Fig 1A, C) in the lungs of 5 of 6 animals at day 3 post-infection, which resolved by day 9. Total genomic Nucleocapsid (gN) and subgenomic Nucleocapsid (sgN) RNA levels from nasal and throat swabs peaked 1 to 2 days post-infection and decreased to undetectable levels by day 7 to 10 (Fig 1D). Viral RNA was also found in the BAL of all animals at day 4 post-infection and was mostly cleared by day 7-10. It should be noted that day 4 post-infection likely does not represent the peak of viremia in the BAL, and previous studies indicate peak viral loads are reached at day 1 post infection in the BAL^44^. Viral RNA was essentially absent from plasma at all timepoints, consistent with previous reports^44^.

**Figure 1:**
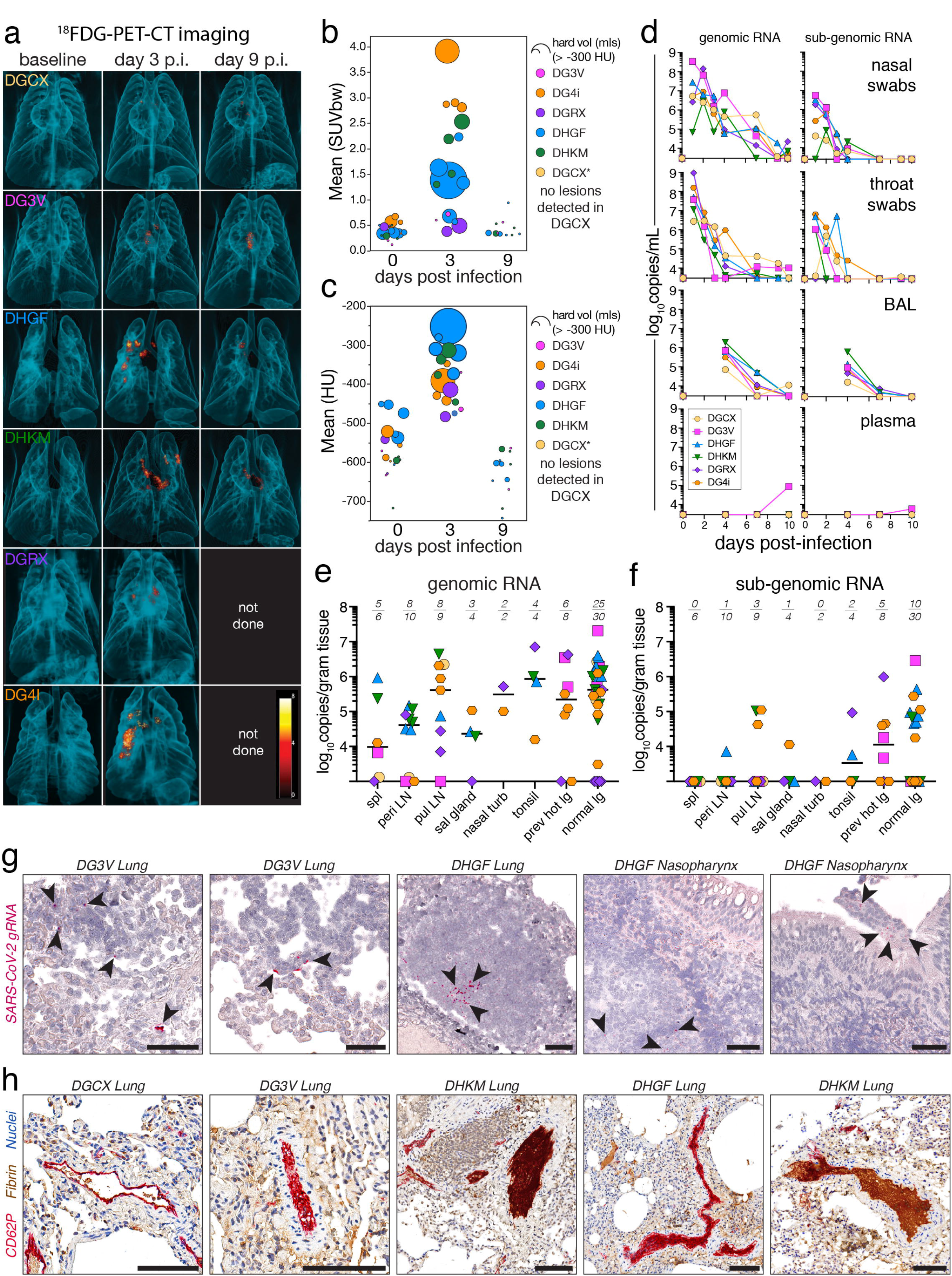
Mild disease and rapid viral clearance in rhesus macaques infected with SARS-CoV-2. Six rhesus macaques infected with 2x10^6^ TCID_50_ of SARS-CoV-2/WA-1 intranasally (1x10^6^) and intratracheally (1x10^6^). (A) 3D rendering of lung ^18^FDG-PET/CT images pre-infection, day 3 and 9 post-infection. (B) Quantification of the metabolic activity (mean ^18^FDG SUV) and volume of tissue with >-300 Houndsfield units (HU) (size of dot) from individual lesions, based on VOI defined at day 3 post infection. (C) Quantification of density (mean HU) and volume of tissue with > -300 HU (size of dot) from individual lesions, based on VOI defined day 3 post infection. DGCX did not have any detectable lung lesions. DGRX and DG4i did not have PET/CT imaging done at day 9 post infection. (D) Quantification of viral genomic RNA (left column) and subgenomic RNA (right column) of the N gene from nasal swabs, throat swabs, BAL, and plasma in copies/mL by RT-qPCR. Cutoff for positivity for genomic RNA is 3000 copies/mL, cutoff for subgenomic RNA is 2500 copies/mL (nasal/throat) or 3000 copies/mL (BAL/plasma). (E-F) Quantification of viral genomic RNA (E) and subgenomic RNA (F) of the N gene from tissues at day 10 post infection in copies/gram of tissue by RT-qPCR with individual samples and median. Numbers above graph indicate number of samples with positive values (numerator) over total number of samples tested (denominator). Cutoff for genomic RNA is CT>35 and 1000 copies/gram tissues, cutoff for subgenomic RNA is CT >37 and 1000 copies/gram tissue. (G) Representative images of staining for SARS-CoV-2 genomic RNA by RNA scope from the lung and nasopharynx at day 10 post infection. Red is viral RNA. (H) Representative images of clotting patterns in the lung at day 10 post infection. Red is platelet staining for CD62P, brown is fibrin, and blue are nuclei.

**Table 1:** Study animal information.

| animal ID | sex | Age<br>(years) | weight<br>(kg) | experimental<br>round |
| --- | --- | --- | --- | --- |
| DGCX | Male | 5.58 | 7.86 | 1 |
| DG3V | Male | 4.58 | 6.81 | 1 |
| DHGF | Male | 2.58 | 3.7 | 2 |
| DHKM | Male | 2.92 | 3.96 | 2 |
| DGRX | Male | 4.83 | 9.68 | 3 |
| DG4i | Male | 4.83 | 10.01 | 3 |

The animals were necropsied at day 10 post-infection for tissue analysis. A 3D reconstruction of the day 3 PET/CT images with conducting airways was used to locate and individually collect the previously PET hot lung regions and normal lung tissue separately. SARS-CoV-2 gN RNA was found on day 10 in all secondary lymphoid organs (SLO) and non-lymphoid tissues (NLT) tested, including the previously PET hot and normal lung tissue, nasal turbinates, salivary gland, and tonsils (Fig 1E). sgN RNA was present at lower levels compared to gN RNA, and was highest in lung tissue (Fig 1F). The persistence of viral RNA at day 10, was confirmed with RNA scope immunohistochemical analysis (Fig 1G). There was a correlation between genomic and subgenomic RNA levels in the mucosal swabs, BAL, and tissues with detectable RNA (Fig S1A-C). We did not observe a correlation between lung lesion severity at day 3 and viral RNA levels from nasal swabs and BAL at day 1 and 4, respectively (Fig S1D-G). Consistent with previous reports in macaques, various forms of microthrombi were still detectable on day 10 post-infection (Fig 1H)^45^. Thus, in rhesus macaques SARS-CoV-2 viral loads peak ∼1-2 days after exposure and this results in mild and transient radiographic evidence of lung inflammation at ∼3 days post-infection, with residual viral RNA in tissues and microthrombi in the lungs at day 10.

### Longitudinal scRNAseq analysis of BAL and PBMC

To compare cellular immune responses in circulation versus airways, single cell RNA sequencing was performed on cryopreserved peripheral blood mononuclear cells (PBMC) and BAL samples obtained prior to infection and at days 4, 7 and 10 post-infection. Uniform manifold approximation and projection (UMAP) and nearest neighbor clustering of PBMCs from all timepoints identified multiple myeloid and T/NK cell populations along with B cells, platelets, and a mixture of proliferating cells (Fig 2A). Due to PBMC isolation and cryopreservation, granulocyte populations were not accounted for in this study. Myeloid and T/NK cell populations were selected for subsequent clustering. We identified nine distinct T/NK cell subsets in PBMCs across all timepoints (Fig S2). Overall, we did not detect major alterations in the T/NK cell composition from PBMCs, but at day 4 after infection, we did observe a drop in naïve CD8 T cells and an increase in central memory CD4^+^ T cells (PBMC T/NK subpopulation 0 and 1, respectively) (Fig S2). Further clustering of myeloid cells identified seven distinct myeloid subsets in PBMCs (Fig 2B-C). Most strikingly, there were major changes to CD14^+^ monocytes after infection. At baseline, a subpopulation of CD14^+^ monocytes expressing *PGTS2* (PBMC myeloid subpopulation 3) were predominant (Fig 2C-D). At day 4 post-infection, there was a dramatic loss of the PGTS2^+^ monocytes with an accompanying increase in two inflammatory monocyte populations with IFN responsive gene signatures (PBMC myeloid subpopulation 0 and 1) (Fig 2C-F). PBMC myeloid population 1 had a more prominent expression pattern of IFN stimulated genes at day 4 as compared to PBMC myeloid population 0, i.e., *MX1*, *MX2*, *IFI6*, *IFI16*, *IFI27*, *ISG15*, and *OAS2*, although both populations showed evidence of response to IFN (Fig 2B,E-F). In contrast to the major changes in CD14^+^ monocytes, CD16^+^ monocytes (PBMC myeloid population 5) did not increase in relative abundance after infection (Fig 2B-D).

**Figure 2:**
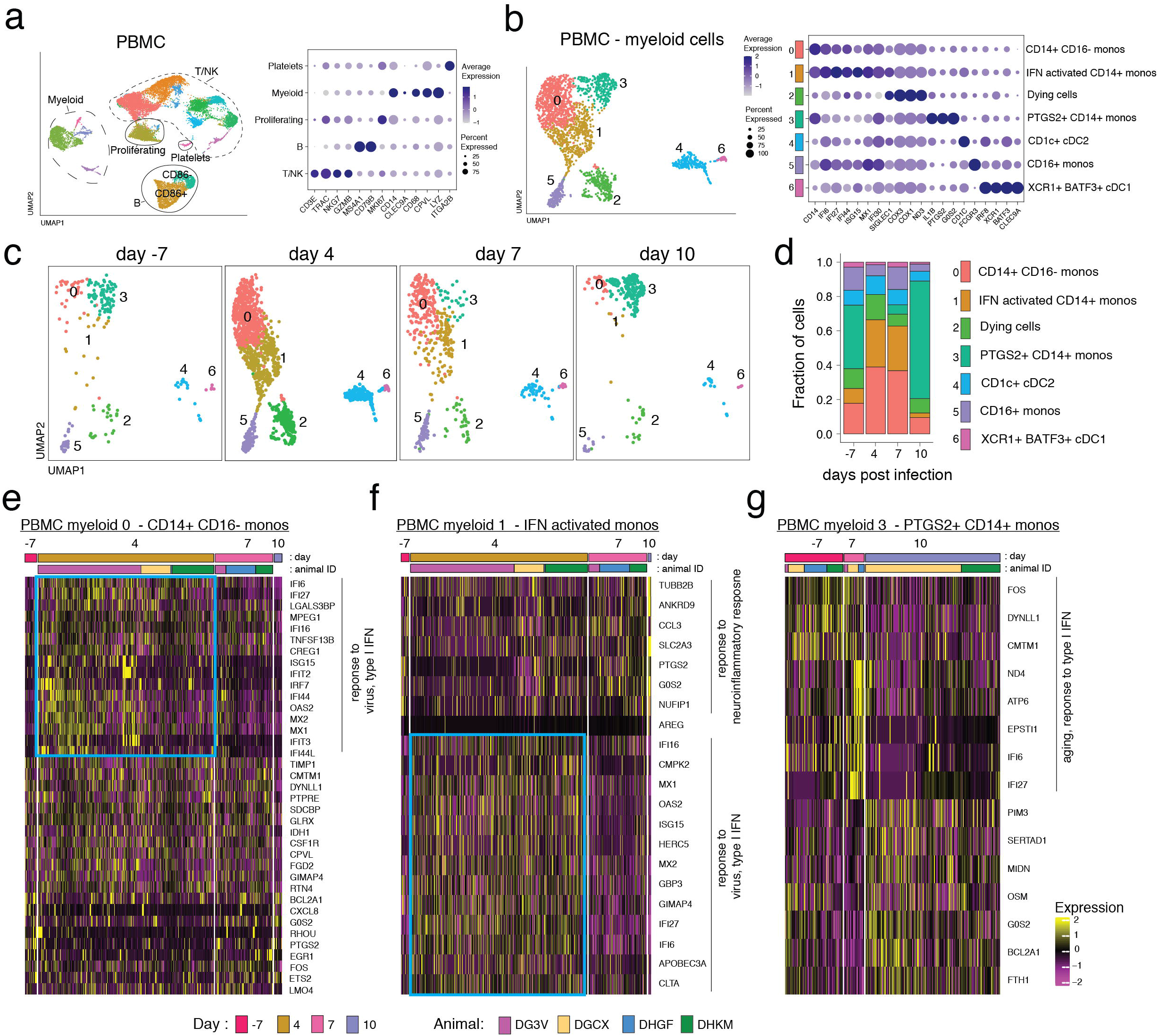
Rapid and transient alterations in CD14^+^ monocytes in PBMCs after SARS-CoV-2 infection. (A) UMAP plot representing the clustering pattern of cells from scRNA-seq data of PBMCs from 4 animals (DGCX, DG3V, DHGF, and DHKM) (left panel). Each dot denotes a cell and is colored based on automated cluster identification. Clusters of cells belonging to a certain cell-type are demarcated and indicated on the plot. Expression levels of cell type defining markers are shown as a dot plot (right panel). Color intensity and dot size represent level of expression and percent of cells in that cluster expressing the gene as defined in the key. (B) UMAP representation of the sub-clustering of the myeloid cells from *A*. Clusters were annotated with cell-types based on gene expression patterns as shown on the dot plot and are identified with different numbers and colors on the plots. (C) UMAP plots separated by time depict the kinetic of the myeloid cells characterized in *B* at pre-infection (d-7), and day 4, 7, and 10 post-infection. (D) Fraction of cells that comprise each myeloid cell-type for each of the 4 timepoints shown in *C* is summarized. (E-G). Heatmap represents the hierarchical clustering of normalized expression levels of differentially expressed genes for each cell for three myeloid clusters. The cluster names are indicated on top of the heatmap and the first and second color bars distinguish time point and animal respectively. Genes were considered differentially expressed between timepoints if log fold change ≥ 0.5 and adjusted p-value < 0.01. Biological processes associated with the genes are indicated on the side and the blue box highlights type I IFN responsive genes upregulated at day 4.

An increase in certain subsets of dendritic cell (DC)2 have been associated with moderate/severe disease in COVID-19 patients^66, 67^. At day 4 post-infection we observed an increase in CD1c^+^ conventional DC2s (cDC2^68^) (PBMC myeloid subpopulation 4), which contracted by day 10. Conventional DC1 cells (*XCR1*, *BATF3* -expressing PBMC myeloid population 6) were less abundant than cDC2s and changed relatively little in abundance during infection. The major alterations in the CD14^+^ monocytes substantially declined by day 7 post-infection and returned to baseline levels by day 10 (Fig 2C-D). The *PGTS2*-expressing monocytes that were lost at day 4 returned by day 10 post-infection, and did not show dramatic changes in gene expression (Fig 2C-D,G).

In the BAL multiple distinct T cell and myeloid populations were identified, along with proliferating cells, B cells, plasmacytoid DCs (pDC), MAST cells, and epithelial cells (Fig 3A). Further clustering of BAL T cells identified 5 populations of T cells (Fig S2D). The largest change was the appearance on day 4 of a population with a mixture of CD8 and CD4 T cells that had a prominent IFN-stimulated gene signature (BAL T cell sub population 3) (Fig S2D-F). These IFN-activated T cells were no longer detectable by day 7 post-infection. Further clustering of BAL myeloid cells revealed 10 distinct populations of myeloid cells (Fig 3B-C). At baseline, BAL cells were mostly comprised of multiple MRC1^+^MARCO^+^ myeloid subsets (BAL myeloid subpopulations 0, 2, and 3) which are likely alveolar macrophages (Fig 3C). At day 4 post infection, there were major increases in populations of IFN-activated monocytes and macrophages in the BAL (BAL myeloid subpopulation 1 and 6), which declined by day 7 and returned to baseline levels by day 10 (Fig 3B-D). At day 10 post-infection, the myeloid cells in the BAL were dominated by a population of CD1c^+^ cDC2s (BAL myeloid sub population 4) (Fig 3C). The cDC2 in the BAL had a pattern of differentially expressed genes that suggested that this population also responded to infection by upregulating type I IFN responsive genes at day 4 post-infection (Fig 3E). By day 10 the cDC2 had down regulated the type I IFN genes and upregulated genes associated with responses to lipopolysaccharide (LPS), including additional chemokines and *IL1B*, as well as the macrophage markers *MRC1* and *MARCO*.

**Figure 3:**
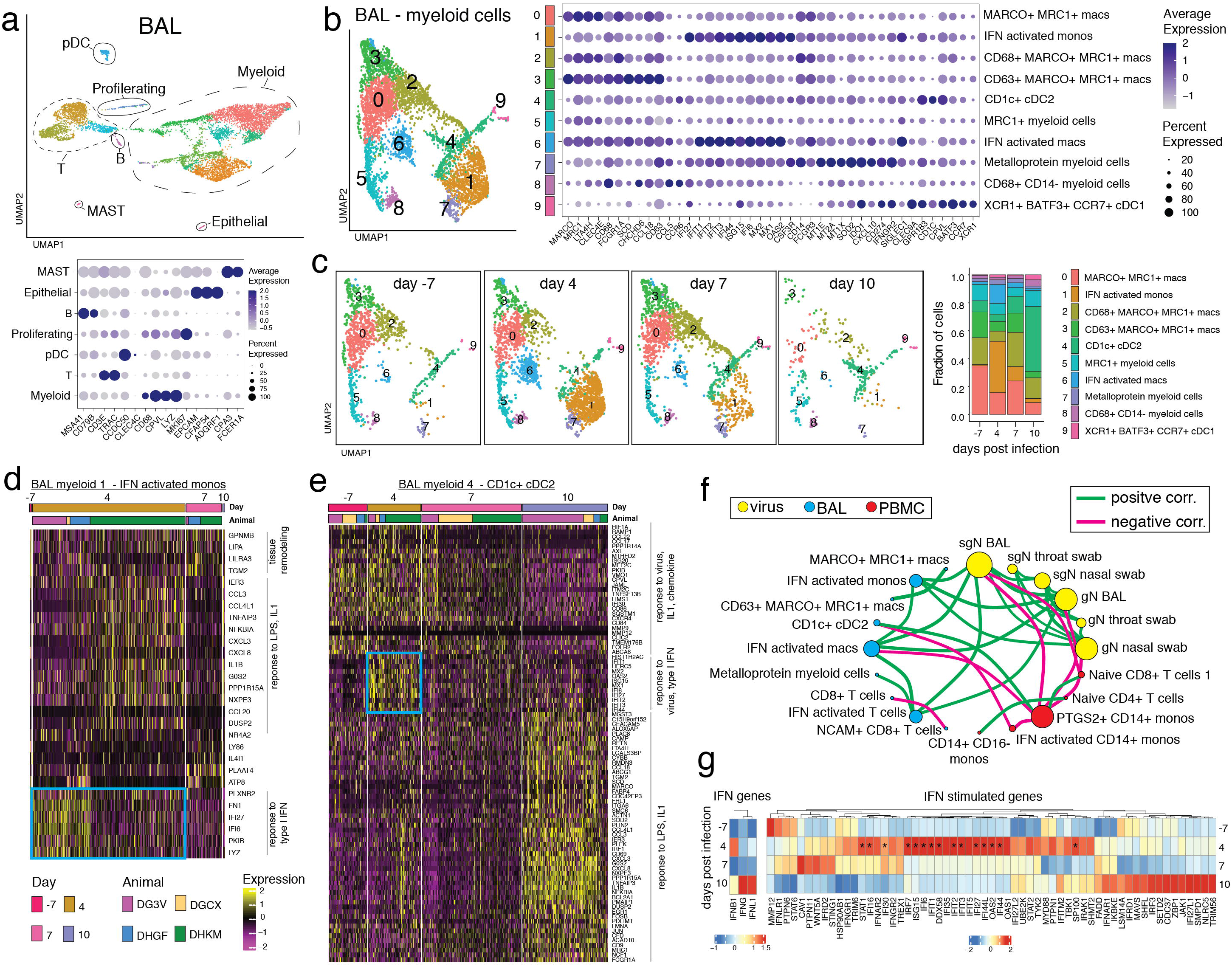
Myeloid cell activation in the airways after SARS-CoV-2 infection. (A) UMAP plot of scRNAseq data from BAL of 4 rhesus macaques (DGCX, DG3V, DHGF, and DHKM) (top panel). Cell clusters are annotated based on broad cell-types and are circled and indicated on the plot. Each dot represents a cell and is colored by cluster. Dot plot displays expression level of markers used to identify the cell types (lower panel). Color intensity and dot size represent level of expression and percent of cells in that cluster expressing the gene marker as defined in the key. (B) UMAP plot (left) of the sub-clustering of the myeloid cells from *A*. Clusters were annotated with cell-types based on gene expression patterns as shown on the dot plot and are identified with different numbers and colors on the plots. (right). (C) UMAP plots depict the kinetic of myeloid cells over time (left) and the fraction of cells that compromise each cluster at pre-infection (d-7), and day 4, 7, and 10 post-infection is summarized as the bar plot (right). (D-E) Normalized gene expression from cells of two BAL myeloid clusters is visualized as a hierarchically clustered heatmap. The timepoints and animals are indicated as colored bars above the heatmap and are defined in the color key. Only genes that were differentially expressed between timepoints (log fold change ≥ 0.5 and adjusted p-value < 0.01) are shown. Biological processes associated with the genes are indicated on the side and the blue box highlights type I IFN responsive genes upregulated at day 4. (F) Spearman’s correlation matrix based on the kinetics of viral loads and fraction of cells from BAL and PBMC myeloid and lymphoid clusters was calculated and visualized as a correlation network. Each circle represents a parameter with the different colors indicating a viral, BAL or PBMC cluster parameter. The size of the circle is proportional to the number of significant correlations (adjusted p < 0.05). A connecting line between two parameters indicates a significant correlation with green and pink lines signifying a positive and negative correlation respectively. (G) Average expression of IFN and IFN stimulated genes from all BAL cells separated by time is clustered and represented as a heatmap. Genes that show a significant difference (adjusted p < 0.05) in expression over time are indicated with ‘*’.

Correlation analysis revealed strong positive correlations between viral RNA levels in the BAL, nasal swabs, and throat swabs (Fig 3F). Viral RNA from BAL and nasal swabs was positively correlated with IFN-activated monocytes, macrophages, and T cells in the BAL. In contrast, PGTS2^+^ monocytes and naïve CD8 T cells from PBMCs negatively correlated with viral RNA from nasal swabs and BAL. To ask if type I, II, or III IFN was the stimulus for the IFN gene signature observed in many cell subsets, we analyzed *IFNB1*, *IFNG*, and *IFNL1* gene expression across all cell types (Fig 3G). We found that *IFNB1* was upregulated at day 4 post-infection, the timepoint when IFN-activated immune cells were highest. Interestingly, *IFNG* and *IFNFL1* showed a relative increase at day 10, when viral RNA had already decreased substantially. Across all cell types the most highly upregulated, statistically-significant IFN stimulated genes were those downstream of type I IFN signaling and showed a pattern of upregulation at day 4 post-infection. Together, these data indicated that SARS-CoV-2 infection induces a robust type I IFN-activated myeloid cell response in PBMC and BAL, which coincides with radiographic indications of inflammation and resolves along with viral RNA levels between day 7 to 10 post-infection.

### Early B cell responses to SARS-CoV-2 infection

We measured multiple B cell subsets in PBMCs and BAL by flow cytometry, including resting naïve B cells (CD20^+^IgD^+^CD95^-^), activated naïve B cells (CD20^+^IgD^+^CD95^+^), germinal center B cells (GC B cells: CD20^+^IgD^-^BCL6^+^Ki67^+^), plasmablasts (CD20^+^IgD-BCL-6^-^CD38^hi^CD27^+^), and activated memory B cells (CD20^+^IgD^-^BCL-6^-^CD95^+^) (Figs S3A-B). Activated memory B cells were further subdivided into IgM^+^, IgG^+^, IgA^+^, and isotype undefined. After infection, we observed a decrease in total B cells in PBMCs (Fig S3C), an increase of 2-3% in the proportion of activated naïve B cells from PBMC at day 4 and 7 (Fig S3D), and a decrease in the overall proportion of activated B cells in PBMCs that are isotype undefined (Fig S3E). At necropsy, the frequency of B cells varied across tissues. While the spleen had the largest fraction of B cells, the BAL and lung had the highest proportion of activated memory B cells (Fig S3D-G). Anti-spike IgM and IgG were detectable in the plasma and BAL at day 10 post-infection in most animals, although levels were only <2 fold above background (Fig S3I-K). Overall, we detected very few changes in B cell populations, and antibody responses were just becoming detectable by day 10 post-infection.

### Kinetics of SARS-CoV-2-specific effector CD8 and CD4 T cell responses in the BAL and PBMC

We next performed a flow cytometric analysis of the Ag-specific T cell response to SARS-CoV-2. We observed only minor changes in the activation of bulk T cell responses in the PBMC after infection, with more dynamic changes in the BAL after infection (Fig S4A-B). To examine SARS-CoV-2-specific T cell responses, we performed intracellular cytokine staining after *ex vivo* restimulation with peptide pools from the viral spike (S), nucleocapsid (N), and membrane (M) proteins, as well as peptide pools (megapools) derived from multiple SARS-CoV-2 antigens found to be immunogenic in humans^69, 70^. As expected, Ag-specific T cell responses were not detected at day 4 post-infection in PBMCs or BAL (Fig4A-D). CD4 T cell responses to S, N, and megapool, each reached ∼4-6% on average by day 7, whereas Ag-specific CD8 T cells were ∼1% at this timepoint in the BAL(Fig 4B,D). Consistent with a slightly delayed response, Ag-specific CD8 T cells in the BAL continued to expand in frequency and maintained Ki67 expression between days 7 and 10 post-infection, while Ag-specific CD4 T cells peaked in frequency at day 7 and decreased Ki67 expression between days 7 and 10 (Fig 4D-E). Of note, frequencies of Ag-specific T cells were ∼10 to 20-fold higher in the BAL vs. PBMC. Moreover, CD8 and CD4 T cell responses against S and N were consistently immunodominant in comparison to M-specific T cells.

**Figure 4.**
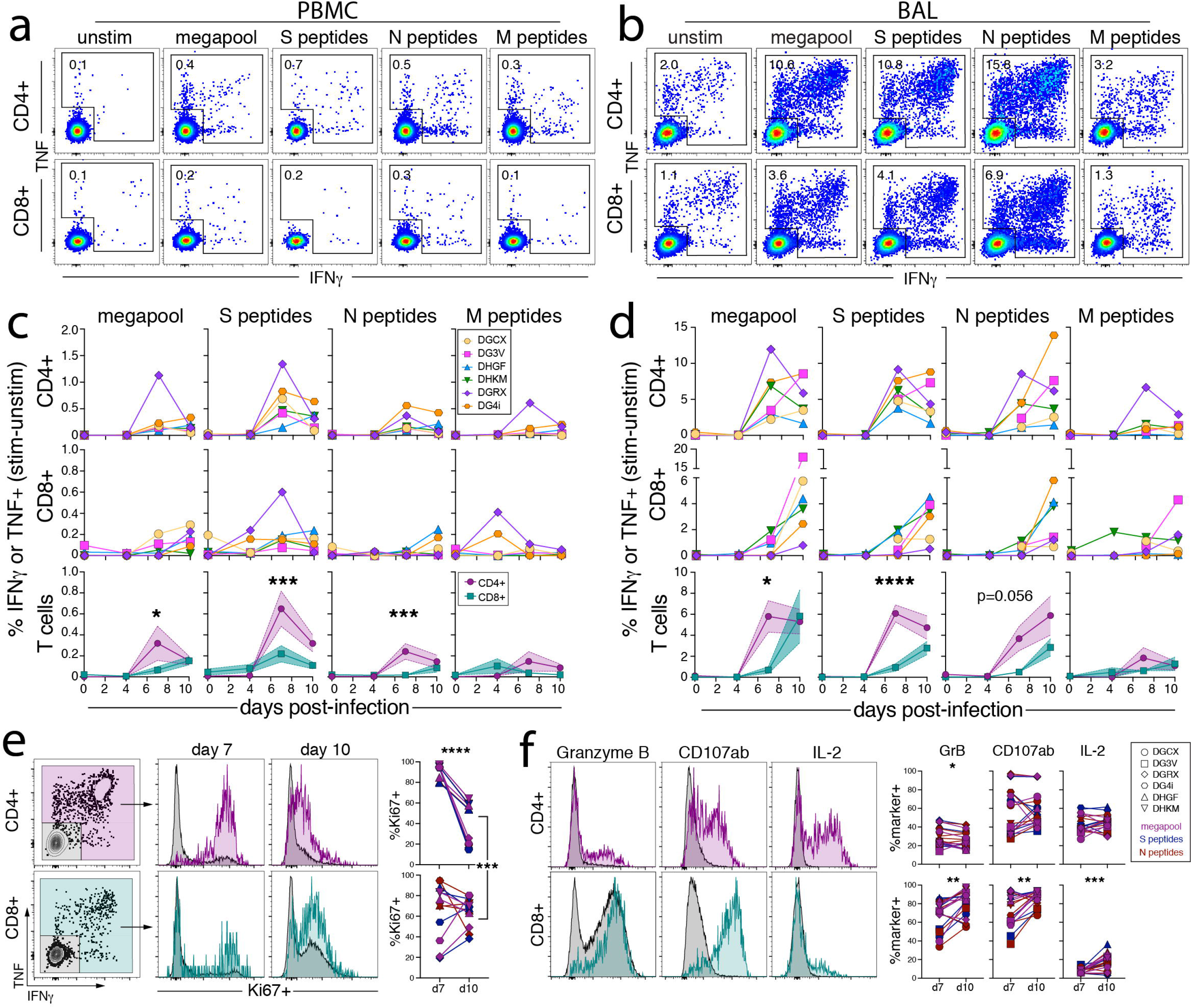
Kinetics of SARS-COV-2-specific CD8 and CD4 T cell responses in the airways. (A-D) Ag-specific CD8 and CD4 T cell responses in the blood and BAL enumerated by production of cytokines (IFNγ and/or TNF) after *ex vivo* peptide stimulation with peptide pools to Spike (S), Nucleocapsid (N), Membrane (M), and an optimized SARS-CoV-2 peptide megapool. Representative flow cytometry plots of Ag-specific CD8 and CD4 T cells from ID#DG4i at day 10 post infection from unstimulated, megapool, S, N, and M peptides from blood (A) and BAL (B), gated on activated T cells i.e., CD8^+^CD95^+^ or CD4^+^CD95^+^. Quantification of Ag-specific T cells from all animals over time in blood (C) and BAL (D), calculated by subtracting the frequency of IFNγ^+^ and/or TNF^+^ in the unstimulated samples from the frequency in the stimulated samples. Bottom row of graphs is an overlay of the mean CD8 and CD4 Ag-specific responses with standard error and a Dunnett’s multiple comparison test of CD4 vs. CD8 responses at each timepoint, p-value <0.05 is considered significant. DGCX and DG3V do not have quantification from day 4 BAL of S, N, and M responses, and are only represented by megapool at day 4. (E) Representative flow cytometry plots of Ki67 expression by Ag-specific CD8 and CD4 T cells from the BAL after S peptide stimulation at day 7 and day 10 post infection from ID#DG4i. Graphs indicate the percent Ki67^+^ of Ag-specific CD8 and CD4 T cells responding to megapool, S, and N peptides from BAL at day 7 and day 10 post infection. Only samples with >35 data points were included. Paired t-test of day 7 vs. day 10 for CD8 and CD4 separately, and CD8 day 10 vs. CD4 day 10. Ki67 staining was not done for ID#DGCX and DG3V. (F) Representative flow cytometry plots of Granzyme B, CD107a/b, and IL-2 expression by Ag-specific CD8 and CD4 T cells from the BAL after S peptide stimulation at day 7 and day 10 post infection from ID#DG4i. Graphs indicate the percent Granzyme B^+^, CD107a/b^+^, or IL-2^+^ of Ag-specific CD8 and CD4 T cells responding to megapool, S, and N peptides from BAL at day 7 and day 10 post infection. Only samples with >35 data points were included. Paired t-test of day 7 vs. day 10 for CD8 and CD4 separately.

In addition to producing IFNγ and TNF after peptide stimulation, the majority of Ag-specific CD8 T cells in the BAL also expressed granzyme B and degranulated after restimulation, as indicated by CD107a/b surface staining (Fig 4F). Approximately 25-60% of Ag-specific CD4 T cells in the BAL also made IL-2. Furthermore, both CD8 and CD4 Ag-specific T cells in the BAL upregulated markers of tissue residence, CD69 and CD103, between days 7 and 10 post infection (Fig S4C-D). Thus, SARS-CoV-2-specific CD8 and CD4 T cells in the airways displayed typical effector functions associated with CTL and Th1 cells, respectively.

### Distribution of SARS-CoV-2-specific CD8 and CD4 T cell responses in mucosal tissues

At the day 10 necropsy, we examined SLO and NLT from the upper and lower respiratory tract for bulk and Ag-specific T cells. Tissue resident memory CD8 and CD4 T cells (CD95^+^CD69^+^CD103^+/-^) were detected in all nonlymphoid tissues measured, including the lung, nasal turbinates, salivary glands, and tonsils (Fig 5A-B). CD103^+^ Trm were more abundant among CD8 compared to CD4 T cells in the BAL, salivary glands, and lymph nodes, which has been shown in other model systems^71, 72^. Using intravenous (i.v.) antibody staining to distinguish between tissue parenchymal and intravascular cells^73–75^, we confirmed that most cells in the BAL, nasal turbinates, salivary gland, tonsils, and lymph nodes were from the tissue parenchyma (Fig S5). As expected for such a highly vascularized tissue, most cells from the lung tissue were intravascular stain positive, but a small population of CD69^+^iv^-^ cells were detectable in the lungs confirming that tissue resident cells were also detected in pulmonary tissue. We next quantified the magnitude of SARS-CoV-2-specific T cells in each of these tissues. S, N, and megapool-specific CD8 and CD4 T cells were detected in the BAL, previously PET hot lung lesions, pulmonary lymph nodes, peripheral lymph nodes, spleen, and PBMC (i.e. the frequency of IFNγ^+^ and/or TNF^+^ cells after peptide restimulation was statistically significantly higher than the unstimulated samples). Surprisingly, Ag-specific CD8 and CD4 T cell responses could not be detected in the majority of nasal turbinates, salivary gland, and tonsils (Fig 5C). The absence of Ag-specific T cells cannot be accounted for by poor T cell isolation from tissues (Figure 5A-B) or lack of virus replication at these sites (Fig 1D-F). Thus, the early clonal burst of SARS-CoV-2-specific T cells is highly skewed toward the BAL and unexpectedly undetectable in the nasal and oral mucosa.

**Figure 5.**
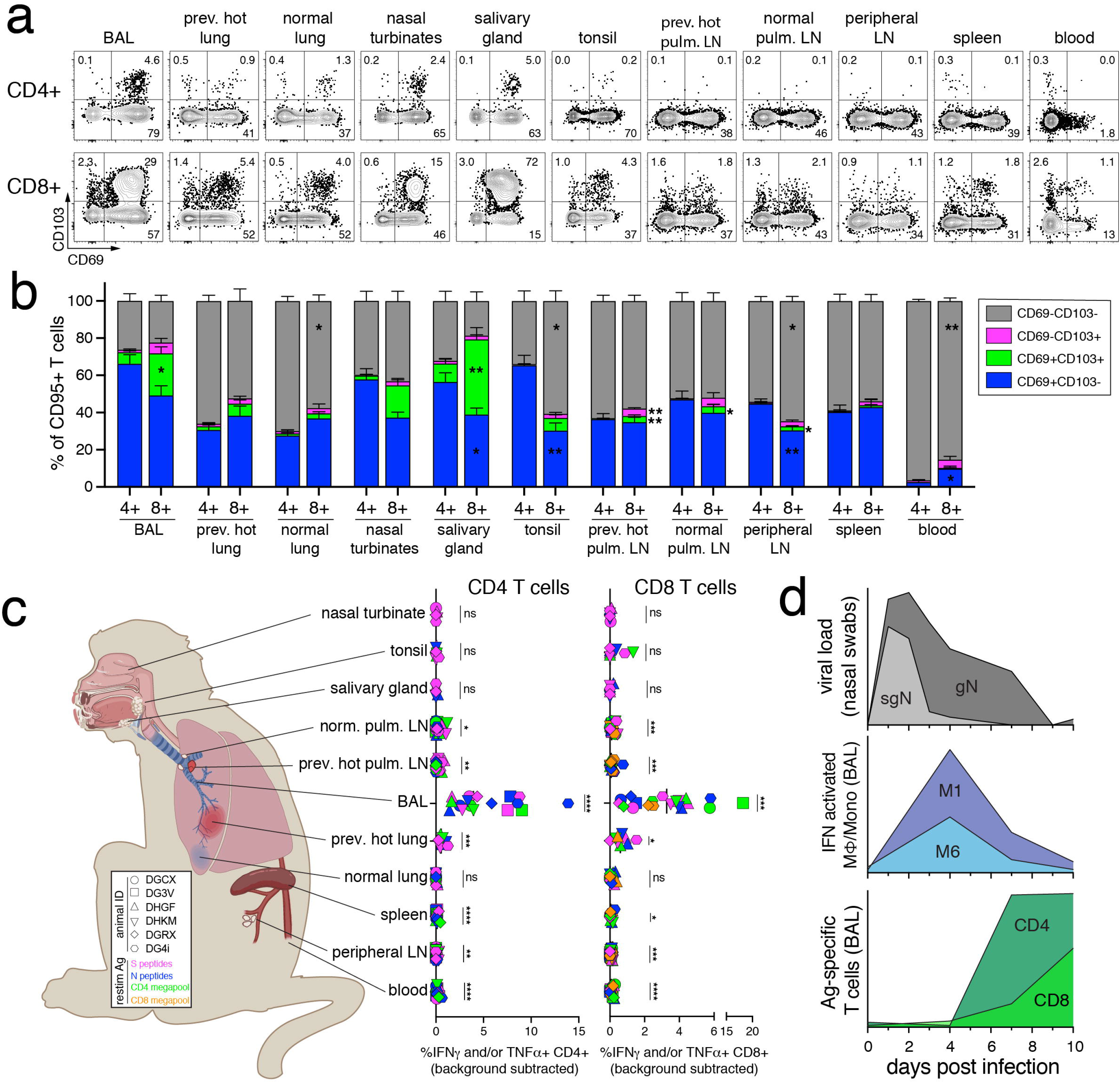
Distribution of SARS-CoV-2-specific effector CD8 and CD4 T cells in mucosal tissues. (A-C) CD69 and CD103, and antigen-specific T cell responses after *ex vivo* peptide stimulation from secondary lymphoid organs and non-lymphoid tissues at day 10 post infection. (A) Representative flow cytometry plots of CD69 and CD103 expression on CD8^+^CD95^+^ or CD4^+^CD95^+^, from BAL, previously hot lung sections, normal lung sections, nasal turbinates, salivary gland (parotid), tonsil, previously hot pulmonary lymph node, normal pulmonary lymph node, peripheral lymph node (axillary), spleen, and blood from ID#DHGF in unstimulated samples at day 10 post infection. (B) Enumeration of percent CD69^+^ and CD103^+^ of CD8^+^CD95^+^ or CD4^+^CD95^+^ in unstimulated samples. Peripheral lymph node includes axillary, inguinal, and cervical lymph nodes. Previously hot lung sections not done for DGCX and DG3V. Sidak’s multiple comparison test for values on CD8 vs. CD4 T cells for each tissue. (C) Tissue distribution diagram and quantification of the frequency of Ag-specific (IFNγ^+^ and/or TNF^+^) in CD4^+^CD95^+^ (left graph) or CD8^+^CD95^+^ (right graph) in each tissue. Frequency calculated by subtracting the frequency of IFNγ^+^ and/or TNF^+^ in the unstimulated samples from the frequency in the stimulated samples. Animals are indicated by shapes and stimulation peptide by color. Statistics are paired t-tests of stimulated vs. unstimulated for each condition. Raw values not plotted. Ag-specific T cells from BAL and blood at day 10, was shown in figure 4, with addition of CD8 megapool here. Tissue graphic created with BioRender.com (D) Representative summary graphs of the immune response to SARS-CoV-2 infection in rhesus macaques. Top graph: median genomic and subgenomic viral RNA levels from nasal swabs on a log scale, as in figure 1. Middle graph: mean frequency of myeloid subpopulation 1 and 6 in BAL, as in figure 3. Bottom graph: mean frequency of the sum of Ag-specific T cells (S+N+M peptide pools) in BAL, as in figure 4.

Overall, the kinetics of SARS-CoV-2 replication and innate/adaptive immune response in rhesus macaques appears typical of an acute viral infection (Fig 5D). SARS-CoV-2 replication peaks within 1 to 2 days post-infection and rapidly decreases thereafter. IFN-responsive myeloid responses are rapidly detected in the PBMC and BAL at day 4 post-infection. Innate immune responses and lung inflammation decline by day 7 post-infection, as Ag-specific T cells begin to accumulate in the airways.

## DISCUSSION

We show here that during mild COVID-19 in rhesus macaques, SARS-CoV-2 replication is largely suppressed prior to the induction of virus-specific T cell responses. PET/CT imaging showed regions of ground glass opacity and consolidation with elevated ^18^FDG uptake in the lungs on day 3 after SARS-CoV-2 infection, which completely resolved by day 9. A longitudinal scRNAseq analysis identified early type I IFN responsive monocyte, macrophage, and dendritic cells in PBMC and BAL that mostly dissipated prior to the arrival of virus-specific CD8 and CD4 T cells. SARS-CoV-2-specific effector T cells were abundant in the pulmonary compartment, but undetectable in nasal turbinates, tonsils, and salivary glands, highlighting major differences in localization of antigen-specific T cells into pulmonary and extrapulmonary mucosal tissues during SARS-CoV-2 infection.

Type I IFN is emerging as a critical mediator of control of SARS-CoV-2 infection^5–7^. In our study the abundance of type I IFN activated myeloid cells in the BAL positively correlated with viral loads in the nasal swabs and BAL. These results are consistent with data from African green monkeys by Speranza et al. showing a strong type I IFN gene signature in macrophages from lung tissue three days after SARS-CoV-2 infection^50^. However, several scRNAseq studies from patients with COVID-19 have found that inflammatory monocytes/macrophage populations are increased with disease severity^2, 11, 12, 14^, suggesting that early type I IFN responses are host protective but prolonged activation of this pathway may be detrimental.

Ag-specific T cell responses were substantially greater in the BAL versus PBMCs, with the average sum of spike, nucleocapsid, and membrane-specific T cells reaching ∼12% of CD4 T cells and ∼7% of CD8 T cells in the BAL compared to ∼1% and ∼0.2% in PBMC, respectively. In the BAL, virus-specific Th1 cell responses preceded CTL responses. Indeed, the CD8 T cell clonal burst likely had not yet peaked, evidenced by their maintained expression of Ki67 at day 10 post-infection. The lack of virus-specific T cells in the nasal turbinates, salivary glands, and tonsils, despite virus infection and subsequent clearance from these tissues, was surprising. The mechanisms underlying the lack of antigen-specific effector T cells in the infected nasal and oral mucosa are not clear. It remains possible that SARS-CoV-2-specific T cell responses were not detected in these sites because they produce molecules other than IFNγ, TNF, IL-2, granzyme B, or degranulation markers CD107a/b, after *ex vivo* peptide stimulation. Nevertheless, it is unlikely that T cells in the nasal turbinates, salivary glands, and tonsils have a completely different functional profile compared to their counterparts in the rest of the host. Alternative techniques for functionally agnostic detection of SARS-CoV-2-specific T cell responses, such as the activation induced marker (AIM) assay, should be tested in future studies^17, 70, 76^. There may also be T cells in these tissues specific to antigens other than the ones tested here, although this too seems unlikely, as the peptide pools used contain numerous immunogenic peptides from across the entire viral genome^69^. Lastly, it is possible that T cells accumulate in these tissues after day 10, and further studies will be needed to determine the longevity and breadth of SARS-CoV-2 specific T cell responses in tissues at later time points.

Our findings support the hypothesis that control of primary SARS-CoV-2 infection in these tissues is T cell independent, which is consistent with a report by Hasenkrug et al. showing that rhesus macaques depleted of CD4 and/or CD8 T cells prior to SARS-CoV-2 infection controlled the virus in the upper and lower respiratory tract, albeit perhaps with a slight delay^65^. Another study also found that CD8 depletion in cynomolgus macaques had no impact on control of SARS-CoV-2 infection^77^. It is important to point out, however, that our data do not rule out a critical role for T cells in other settings of SARS-CoV-2 infection. For example, T cells likely play a significant role when SARS-CoV-2 infection does not resolve quickly, such as during moderate and severe COVID-19. T cells have been implicated in control of SARS-CoV-2 in other susceptible animal models, like the human ACE2 expressing mouse lines and Syrian hamsters^78–80^. Furthermore, nucleocapsid specific CD8 T cells are correlated with less severe disease in patients^20^. In addition, preclinical studies suggest that depletion of CD8 T cells from vaccinated monkeys prior to SARS-CoV-2 challenge significantly impairs control of virus replication^60^. T cells may also play a major role in vaccine-elicited protection, and T cell targeted peptide vaccines are currently being developed^20, 81, 82^. Vaccine-elicited T cells may prove critical in protection against SARS-CoV-2 variants of concern that are able to evade neutralizing antibodies, as T cell epitopes are thought to be more conserved across isolates^83^. In our study, it should also be noted that T cells may have played a role in clearance of virus-infected cells remaining in the lungs after the first week, when T cells arrived in the tissue.

Altogether, these data show that mild SARS-CoV-2 infection is associated with effective innate immune-mediated control. Future studies are needed to determine the importance of individual innate and adaptive immune cell types in suppression of SARS-CoV-2 replication.

## MATERIALS AND METHODS

### Study design

All animal experiments were approved by Animal Care and Use Committee (ACUC) and all methods were approved on animal safety protocol LPD-25E at the National Institute of Health. Experiments were conducted in an AAALAC accredited aBSL-3 vivarium facility in Bethesda, Maryland. Animals were singly housed in vented air cages with a 12-hour light/dark cycle. The animals were monitored twice daily, with a detailed physical exam once per day during the study. The Institutional Biosafety Committee approved all work with SARS-CoV-2 in the BSL-3 level facility and approved any inactivation methods used.

### Animals and infection

Six, male rhesus macaques aged 2.5 to 6 years, weighing 3-10 kg were infected with SARS-CoV-2/USA/WA-1 (Table 1). For infection, animals were anesthetized as described below and administered 2x10^6^ TCID_50_ total: 1x10^6^ TCID_50_ in 3mL intratracheally with a plastic gavage tube attached to 5 mL syringe, and 5x10^5^ TCID_50_ in 0.5mL intranasally in each nostril. The animals were examined daily with a health scoring sheet, as previously described^44^. Animals were anesthetized with ketamine and dexmedetomidine at baseline (day -5 to -30), day 0, 1, 2, 3, 4, 7, and 10 (necropsy) for exams, viral load swabs, blood and BAL fluid draws, and complete blood count and C-reactive protein analysis. During anesthesia, animals were weighed and monitored for heart rate, respiratory rate, body temperature, oxygen saturation. Glycopyrrolate and atipamezole were given for recovery from anesthesia.

### 18FDG-PET/CT Acquisition and Data Analysis

Rhesus were sedated and imaged by PET/CT during mechanical ventilation (Hallowell Ventilator Model 2002) at baseline (day -22 to -5), on day 3, and 9 post infection. To reveal metabolic hyperactivity consistent with inflammation, a [^18^F]-FDG dose of 0.5 mCi/kg was given intravenously 1 hour prior to PET imaging. During the uptake time a high-resolution CT scan of the lungs was acquired with a breath hold on a LFER 150 PET/CT scanner (Mediso Inc, Budapest, Hungary) as previously described^84^. The raw CT and PET data were reconstructed using the Nucline software (Mediso, Inc, Budapest, Hungary) to create individual DICOM files that were co-registered using MIM Maestro (v. 7.0, MIM Software Inc, Cleveland, Ohio).

By aligning baseline PET/CT fused images and those taken at day 3 and 9 in MIM Maestro, specific lung regions with abnormal density (> ∼ -550 HU) or metabolic activity (> ∼ 1.5 SUV) were identified as volumes of interest (VOI) or lesions similar to methods used previously, rather than using whole lung volume of interest^85^. For each animal, the lesion VOIs (day 3 in this study) were transferred to the aligned PET/CT images acquired at baseline and the day 9 time point, adjusting for position variations but keeping the same volume. Disease volume was estimated by using two density thresholds: tissues harder than -550 Hounsfield Units (HU) or harder than -300 HU for evaluating change over time. Regarding metabolic activity, PET parameters were estimated using a threshold of > 2 standardized uptake value (SUV). Similar reference VOIs were used to identify metabolically activated tissues (SUV > 2) within peri-carinal lymph nodes (LN). LN [^18^F]-FDG uptake was measured in activated regions of the hilar and subcarinal LNs of each animal. Our analysis also included calculations of total lesion glycolysis (TLG). Two readers independently performed image analysis for each animal using consistent lesion labeling determined by a third reviewer. Three-dimensional projections of FDG uptake in the lung regions were generated using Osirix v 5.9 software (Pixmeo, Geneva, Switzerland) as previously described^86^.

### Blood and BAL collection

Blood and BAL collection procedures followed ACUC approved standard operating procedures and limits. Blood was collected in EDTA tubes and centrifuged at 2,000rpm for 10 minutes at 22°C to isolate plasma. After plasma removal, remaining blood was diluted 1:1 with 1x PBS. 15mL of 90% Ficoll-Paque density gradient (Cytiva Cat#17144002), diluted with 10x PBS, was added to SepMate™ PBMC Isolation Tubes (StemCell Cat#85450) and centrifuged at 1,000g for 1 minute at 22°C, to collect Ficoll below the separation filter. Blood and PBS mix was added to the SepMate tube with Ficoll-Paque and centrifuged at 1,200g for 10 minutes at 22°C. The upper layer was poured into a 50mL conical and brought to 50mL with PBS +1% FBS, and then centrifuged at 1,600rpm for 5 minutes at 4°C. The cell pellet was resuspended at 2x10^7^ cell/mL in X-VIVO 15 media + 10% FBS for subsequent analysis. BAL was collected after intubation by instillation of 50mL of warm pharmaceutical-grade PBS, 10mLs at a time. For cellular analysis, BAL was filtered through a 100um filter into a 50mL conical and centrifuged at 1,600 rpm for 15 minutes at 4°C. The cell pellet was resuspended at 2x10^7^ cell/mL in X-VIVO 15 media + 10% FBS for subsequent analysis.

### Necropsy

Intravenous antibody was administered prior to euthanasia, as previously described^74^. Briefly, prior to necropsy, 10mL of blood was drawn as a negative control and 100ug/kg of αCD45-biotin (clone: ITS_rhCD45 developed by Roederer Lab) was infused. After infusion the BAL and ∼60mL of blood was collected. After prosection of the lung and airways, specific lung regions observed to have abnormal HU density or FDG uptake in the day 3 images were collected separately from the remainder of the lung. LNs identified as having regions of SUV > 2.5 were collected separately from those with lower SUV on day 3.

### Viral RNA quantification

RNA from the nose and throat was collected by swabbing each nostril or back of the throat, respectively, with a sterile swab for 10 seconds. Swabs were placed in 1mL viral transport media (1x HBSS, 2% FBS, 100ug/mL Gentamicin, and 0.5ug/mL amphotericin B) and stored on ice until RNA extraction. Swabs were vortexed in swab media before removing the swab tip. For RNA extraction 140uL of sample (plasma, 1^st^ BAL wash, or swab media) was processed using a Viral RNA mini kit (Qiagen Cat# 52906) and eluted in 50uL RNase Free water. For RNA isolation from tissues, tissue pieces were weighed before placing in 1mL RNAlater® media (Sigma Cat# R0901) and stored at 4°C overnight and then stored at -80°C long term. Tissues were then thawed and processed in the RNeasy Plus Mini kit (Qiagen # 74136) and eluted in 50uL RNase Free water. Eluted RNA was stored at -80°C long-term.

Extracted RNA was used in a RT-qPCR reaction for detection of total or subgenomic RNA from the N gene of SARS-CoV-2. Total RNA reactions amplify both genomic viral RNA and subgenomic viral mRNAs, and are labeled as genomic throughout the manuscript to differentiate from the subgenomic viral mRNA-specific amplifications. Each sample was prepared in a 12.5uL reaction, with 2.5uL of eluted RNA, 3.25uL Taqpath 1-step RT-qPCR Master Mix (Thermo Cat#A15299), primers at 500nM, probes at 125-200nM, and the remaining volume as RNase free water. N1 genomic RNA was detected with 2019-nCoV RUO Kit, 500 rxn (IDT #10006713), containing CDC 2019-nCoV_N1 Forward Primer (5’-GAC CCC AAA ATC AGC GAA AT-3’), CDC 2019-nCoV_N1 Reverse Primer (5’-TCT GGT TAC TGC CAG TTG AAT CTG-3’), and CDC 2019-nCoV_N1 Probe (5’-[FAM]-ACC CCG CAT TAC GTT TGG TGG ACC-[BHQ1]-3’) at 125nM. N gene subgenomic RNA was detected using Forward Leader sequence primer (5’-CGA TCT CTT GTA GAT CTG TTC TC-3’), sgN Reverse (5′-GGT GAA CCA AGA CGC AGT AT-3’), and sgN Probe (5′-[FAM]-TAA CCA GAA TGG AGA ACG CAG TGG G-[BHQ1]-3′,) at 200nM, all custom made from Eurofins. All samples were tested for RNA integrity using the 2019-nCoV RUO Kit for RNase P, containing CDC RNAse P Forward Primer (5’-AGA TTT GGA CCT GCG AGC G-3’), CDC RNAse P Reverse Primer (5’-GAG CGG CTG TCT CCA CAA GT-3’), and CDC RNAse P Probe (5’-[FAM]-TTC TGA CCT GAA GGC TCT GCG CG-[BHQ]-1-3’). Prepared reactions were read on a QuantStudio 7 Flex Real-Time PCR System, 384-well format (Applied Biosystems Cat# 4485701). Cycling conditions: Initial: 25°C for 2 minutes, 50°C for 15 minutes, and 95°C for 2 minutes, Cycling: 95°C for 3 seconds, 60°C for 30 seconds, for 40 cycles. Copies per/mL or copies/gram were calculated based on standard curves generated for each RT-qPCR run, with RNA standard of known quantity and 10, 5-fold dilutions, run in duplicate. The limit of detection was based on the CT limit of detection from the standard curve in each run. For genomic RNA, this was also limited to CT<35, based on manufacturer’s instructions. For subgenomic RNA cutoff CT<37 was used.

### Tissue digestion

Tissues were processed for single cell suspension before flow cytometry or peptide stimulation as follows. Spleen (approximately 0.5 inch x 0.5 inch portion) and lymph nodes were placed in 5mL of PBS + 1% FBS in a gentleMACS C tube (Miltenyi Cat#130096334) and run on gentleMACS Octo Dissociator (Miltenyi), with m_spleen_02_01 setting, then filtered through a 100um filter into a 50mL conical and centrifuged at 1,600rpm for 5 minutes at 4°C. Salivary gland was gentleMACS dissociated as above, and after centrifugation cells were resuspended in 7mL 44% Percoll® (Sigma Cat# P1644) with 1xPBS and centrifuged at 2,000rpm for 20 minutes at 22°C without brake. The tonsil and lung were gentleMACS dissociated in 5mL digestion buffer (RPMI + 50U/mL DNase I + 1mg/mL hyaluronidase + 1mg/mL collagenase D (Roche)) and then agitated on a shaker at 220rpm for 45 minutes at 37°C. Digestion reaction was stopped with equal parts PBS + 20% FBS and centrifuged at 1,600rpm for 5 minutes at 22°C. The cell pellet was resuspended in Percoll gradient, as above for salivary gland. After processing, the spleen and lung were cleared of red blood cells by resuspending cell pellet in 2mL of ACK Lysing Buffer (Quality Biologicals Cat#118-156-101) for 2 minutes at room temperature, then stopping the reaction with 10-20mL of PBS + 1%FBS. Cells were resuspended at 2x10^7^ cell/mL in X-VIVO 15 media + 10% FBS for further analysis.

### Peptide stimulation assay

Single cell suspensions were plated at 2x10^7^ cell/mL in 200uL in 96 well plates with X-VIVO 15 media, plus 10% FBS, Brefeldin 1000x (Invitrogen Cat#00-4506-51) and Monensin 1000x (Invitrogen Cat#00-4505-51), CD107a APC 1:50, CD107b APC 1:50, and peptide pools at 1ug/mL. Cells were stimulated for 6 hours at 37°C + 5% CO_2_ before surface staining. Spike peptide pool consisted of Peptivator SARS-CoV-2 Prot_S1 (Miltenyi Cat#130-127-048) and Peptivator SARS-CoV-2 Prot_S (Miltenyi Cat#130-127-953). Nucleocapsid peptide pool consisted of Peptivator SARS-CoV-2 Prot_N (Miltenyi Cat# 130-126-699). Membrane peptide pool consisted of Peptivator SARS-CoV-2 Prot_M (Miltenyi Cat# 130-126-703). CD4 megapool consisted of CD4_S_MP and CD4_R_MP, and CD8 megapool consisted of CD8_MP_A and CD8_MP_B, as described^70^. After stimulation cells were centrifuged at 1,600 rpm for 5 minutes at 4°C and proceeded with surface staining.

### Flow cytometry and antibody staining

Cells were resuspended in 50uL surface stain antibodies diluted in PBS + 1% FBS and incubated for 20 minutes at 4°C. Cells were washed 3 times with PBS + 1% FBS, before fixation with eBioscience Intracellular Fixation & Permeabilization Buffer Set (Thermo Cat# 88-8824-00) for 16 hours at 4°C. After fixation cells were centrifuged at 2,200rpm for 5 minutes at 4°C without brake and washed once with eBioscience Permeabilization Buffer. Cells were resuspended in 50uL intracellular stains diluted in eBioscience Permeabilization Buffer, and stained for 30 minutes at 4°C. After staining cells were washed with eBioscience Permeabilization Buffer 2x and resuspended in PBS + 1% FBS + 0.05% Sodium Azide for running on the BD Symphony platform. B cells were resuspended in 50uL Human Fc-Block (BD Cat#564220) diluted to1:500 in PBS + 1%FBS and incubated for 30 minutes at 4°C prior to washing and surface staining.

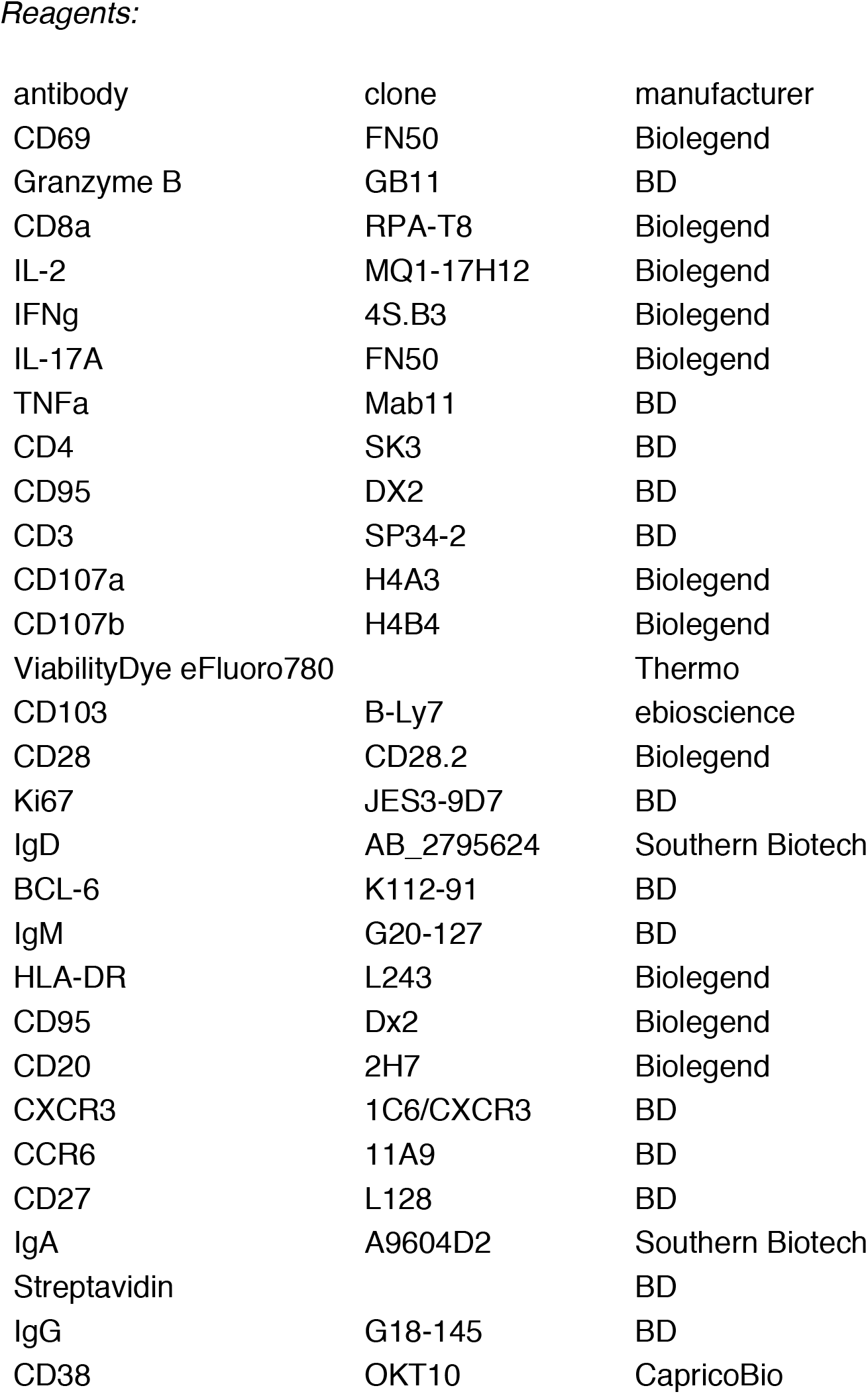
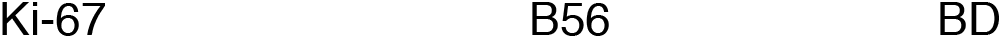

### Immunohistochemistry and RNA scope

Tissue for histology were collected in 10% neutral buffered formalin and stored at room temperature for 16 hours. Fixed tissues were transferred to 70% ethanol and stored at room temperature until processing. Slides were cut 10-microns thick using standard RNAse precautions. Immunohistochemical slides were deparaffinized and treated with AR6 Buffer (Akoya Biosciencecs, USA) for 20 minutes at 100°C. Tissues were then permeabilized using 0.2% TritonX 100 (Millipore Sigma, USA) for 10 minutes. After blocking, slides were incubated with primary antibodies against CD62P (clone EPR22850-190, Abcam, USA) and fibrin (clone 59D8, Millipore Sigma, USA) at a 1:500 and 1:200 concentration, respectively. Following washing, slides were stained according to the protocol for ImmPRESS Duet Double Staining Polymer Kit (Vector Laboratories, USA) and counter-stained with hematoxylin. Slides used for *in situ* hybridization were deparaffinized and treated with RNAscope epitope retrieval buffer (ACD Biotechne, USA) for 20 minutes at 100°C. Endogenous peroxidases were then blocked with hydrogen peroxide and tissue permeabilized with a diluted RNAscope protease plus for 20 minutes at 40°C. Probes for SARS-CoV-2, containing 20 pairs of probes spanning S gene (Category # 848561, ACD Biotechne, USA), were incubated for 2 hours at 40°C. Slides were then processed according to RNAscope 2.5 HD Assay-RED (ACD Biotechne, USA) protocol and counterstained with hematoxylin. Slides stained immunohistochemically or by *in situ* hybridization were imaged using Aperio VERSA (Leica Microsystems, USA) and analyzed using quPath, an open-source software developed by the University of Edinburgh. SARS-CoV-2 puncta were confirmed using both positive and negative controls to ensure accurate staining.

### Single cell RNA sequencing and data analyses

Cells from the BAL fluid and PBMCs from blood were obtained as described above and cryopreserved in 1ml of RPMI + 40% FBS + 15% DMSO. PBMC and BAL samples from days -7, 4, 7, 10 for monkeys DHGF, DG3V and DHKM and both sample types from days -7, 4 and 10 for monkey DGCX were processed for scRNAseq using the 10X Genomics Chromium Single Cell 3’ kit (v3.1). Briefly, cryopreserved samples were quickly thawed using a water bath set to 37°C and washed twice using 10% FBS in RPMI. Samples were then stained with unique TotalSeq-A hashtag antibodies (HTO) as per manufacturer’s (Biolegend) protocol. Equal number of cells from each sample were pooled and super-loaded on a 10X Genomics Next GEM chip and single cell GEMs were generated on a 10X Chromium Controller as previously described^87^. Subsequent steps to generate cDNA and HTO libraries were performed following 10X Genomics and Biolegend’s protocol respectively. Libraries were pooled and sequenced on an Illumina NovaSeq S1 and S2 flow cells as per 10X sequencing recommendations.

The sequenced data was processed using cellranger (version 5.0) to demultiplex the libraries. The reads were aligned to *Macaca mulatta* mmul_10 genome to generate count tables. The count tables were then further processed and analyzed using the Seurat (version 4.0) in R (version 4.1.0). Samples from different PBMC and BAL libraries were integrated using IntegrateData function to account for possible batch effects and to generate one integrated dataset for each tissue type. Cells were then filtered for less than 15% mitochondrial contamination and only singlets as determined by the HTOs were included resulting in 16,769 PMBC and 7,274 BAL cells for downstream analysis. Data were normalized and scaled and FindVariableFeatures function was used to identify variable genes to subset and integrate the data to correct for animal bias. Principal component analysis was performed to find neighbors and clusters and UMAP reduction was performed with 20 dimensions. FindAllMarkers with a filter of log fold change ≥ 0.25 and percent of cells expressing the marker ≥ 0.25 was used to identify gene markers that distinguish the cell clusters, and the clusters were manually assigned cell types based on identified canonical markers. Myeloid and T (and NK in case of PBMC) cell clusters were further subclustered and clusters were again manually annotated based on gene markers determined by the FindAllMarkers function. Differentially expressed genes between timepoints of a particular cluster were identified by running FindMarkers function with MAST and comparing one timepoint to all other timepoints or one timepoint to another in a pairwise manner. Genes with a log fold change ≥ 0.5, percent of cells expressing the marker ≥ 0.25 and adjusted p value ≤ 0.01 were considered significant and these genes were hierarchically clustered and displayed as a heatmap using the ComplexHeatmap function in R. Gene ontology enrichment analysis of genes upregulated at a particular timepoint was performed using clusterProfiler to identify biological processes (adjusted p value ≤ 0.05). The AverageExpression function was used to calculate average gene expression of IFN and IFN stimulated genes across all cells over time and was visualized using pheatmap.

Spearman’s correlation test was performed between viral loads from various sites and fraction of cells in a particular cluster at all available timepoints and filtered for adjusted p value < 0.05. Correlations were visualized using a network diagram generated using igraph in R.

| antibody | clone | manufacturer |
| --- | --- | --- |
| TotalSeq™-A0251 | LNH-94; 2M2 | Biolegend |
| TotalSeq™-A0252 | LNH-94; 2M2 | Biolegend |
| TotalSeq™-A0253 | LNH-94; 2M2 | Biolegend |
| TotalSeq™-A0254 | LNH-94; 2M2 | Biolegend |
| TotalSeq™-A0255 | LNH-94; 2M2 | Biolegend |
| TotalSeq™-A0256 | LNH-94; 2M2 | Biolegend |
| TotalSeq™-A0257 | LNH-94; 2M2 | Biolegend |
| TotalSeq™-A0258 | LNH-94; 2M2 | Biolegend |
| TotalSeq™-A0259 | LNH-94; 2M2 | Biolegend |
| TotalSeq™-A0260 | LNH-94; 2M2 | Biolegend |
| TotalSeq™-A0262 | LNH-94; 2M2 | Biolegend |
| TotalSeq™-A0263 | LNH-94; 2M2 | Biolegend |
| TotalSeq™-A0264 | LNH-94; 2M2 | Biolegend |
| TotalSeq™-A0265 | LNH-94; 2M2 | Biolegend |

## Data availability

Single-cell RNAseq read data will be submitted to NCBI.

## ACKNOWLEDGEMENTS

We would like to acknowledge the Center for Cancer Research Sequencing Facility at the Frederick National Laboratory for Cancer Research for performing the sequencing and Drs. Paul Schaughency and Justin Lack of the NIAID Collaborative Bioinformatics Resource for assistance with the bioinformatics. We would like to thank Dr. Rashida Moore for clinical care of the macaques and Drs. Kerry Hilligan and Paul Baker for assistance with S.O.P.s and inactivation method development for SARS-CoV-2 BSL-3 work. Funding for this study was provided in part by the Division of Intramural Research/NIAID/NIH. A.S. and D.W. were supported by NIH grant contract no. 75N9301900065. The content of this publication does not necessarily reflect the views or policies of DHHS, nor does the mention of trade names, commercial products, or organizations imply endorsement by the U.S. Government.

## AUTHOR CONTRIBUTIONS

C.E.N. led the study. C.E.N., J.M.B., L.E.V., and D.L.B. designed the study. C.E.N., S.N., T.W.F., K.D.K., S.S., D.D., and N.E.L. performed experiments. C.E.N., T.W.F., and D.L.B. analyzed data. S.N., A.S., C.E.N., and D.L.B. performed single cell RNA sequencing analysis. The Tuberculosis Imaging Program (TBIP) managed logistics and performed NHP manipulations including infection, necropsy, PET/CT scanning, and imaging analysis. F.G. and J.D.F. analyzed the PET/CT data. L.E.V. supervised T.B.I.P. and designed the analysis for PET/CT data. C.E.N., S.N., T.W.F, and D.L.B. made figures. E.L.P., M.R., D.W., E.dW., and H.D.H. provided necessary reagents for the completion of the study. C.E.N., S.N. and D.L.B. wrote the manuscript. All authors contributed to editing the manuscript. D.L.B. supervised the study.

---

**Supplemental Figure 1 (refers to Figure 1).**
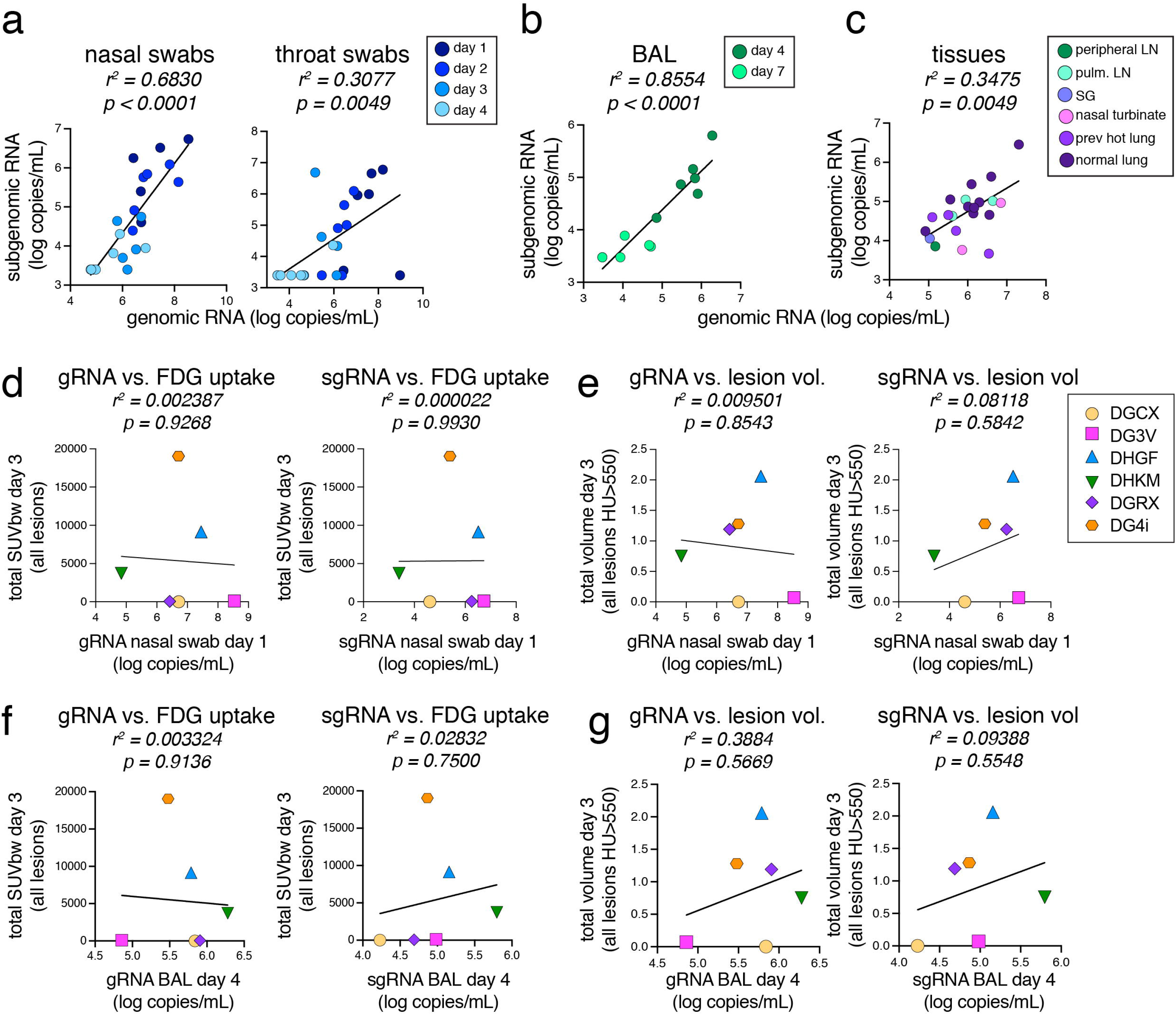
Viral RNA levels are not correlated with disease severity. Correlation analysis of viral RNA and lung lesion severity. (A) Correlation of genomic RNA and subgenomic RNA in copies/mL in nasal swabs (left graph) and throat swabs (right graph) from day 1 to day 4, as shown in figure 1D. (B) Correlation of genomic RNA and subgenomic RNA in copies/mL in BAL from day 4 and day 7, as shown in figure 1D. (C) Correlation of genomic RNA and subgenomic RNA in copies/gram of tissue from day 10 in tissues with values >limit of detection, as shown in figure 1E. (D) Correlation of genomic RNA (left graph) or subgenomic RNA (right graph) at day 1 in nasal swabs and the sum of lung lesion metabolic activity at day 3. (E) Correlation of genomic RNA (left graph) or subgenomic RNA (right graph) at day 1 in the nasal swabs and the sum of lung lesion size at day 3. (F) Correlation of genomic RNA (left graph) or subgenomic RNA (right graph) at day 4 in the BAL and the sum of lung lesion metabolic activity at day 3. (G) Correlation of genomic RNA (left graph) or subgenomic RNA (right graph) at day 4 in the BAL and the sum of lung lesion size at day 3.

**Supplemental Figure 2 (refers to Figure 2 and 3).**
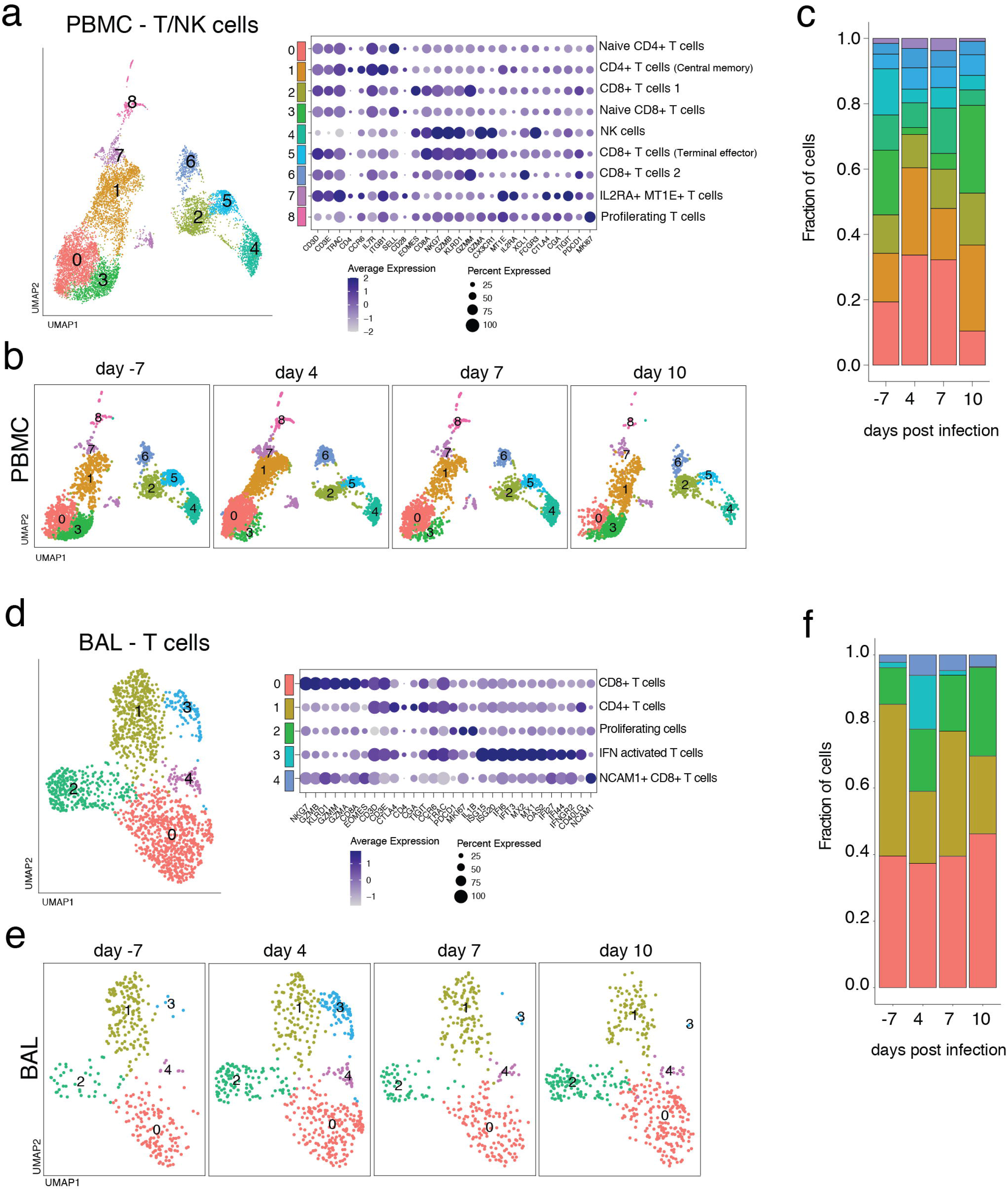
Minor alterations in T cells and NK cells in PBMC and BAL as measured by scRNAseq. (A,D) UMAP plot shows the sub-clustering of PBMC T and NK cells (A) from Figure 2A and BAL T cells (D) from Figure 3A (top panels). Clusters were annotated with cell-types based on gene expression patterns as shown on the dot plot and are identified with different numbers and colors on the plots. (lower panels). (B,E) UMAP plots depict the kinetic of lymphoid cells over time. (C,F) Fraction of cells present in each of the lymphoid cell clusters in PMBC (C) and BAL (F) is summarized.

**Supplemental Figure 3.**
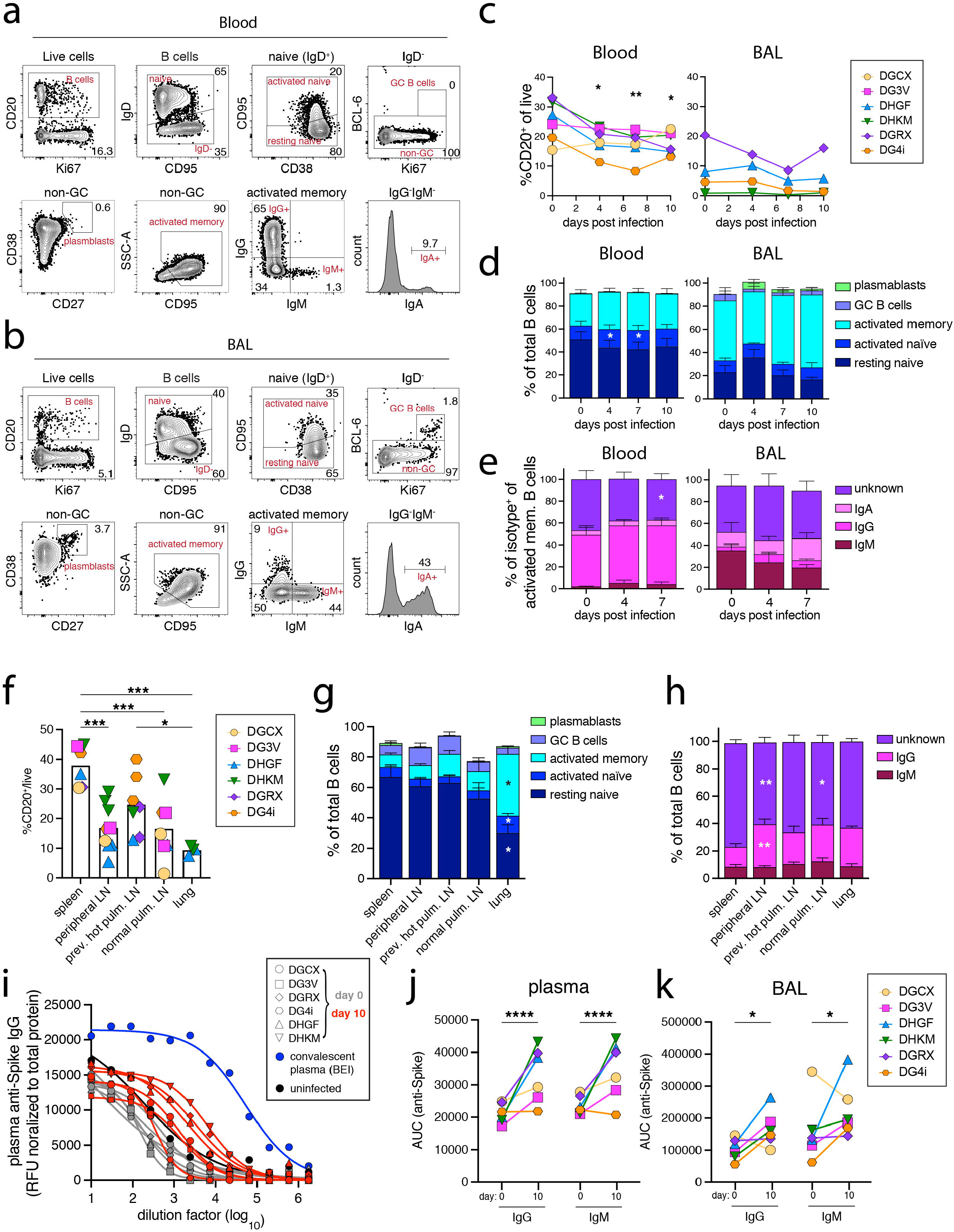
B cell responses after SARS-CoV-2 infection in rhesus macaques. Quantification of different B cell subsets after SARS-CoV-2 infection. Representative flow cytometry gating strategy of B cells and B cell subsets (i.e., resting naïve, activated naïve, germinal center B cells (GC B cells), plasmablasts, activated memory and different isotypes IgG^+^, IgM^+^, and IgA^+^) from the blood (A) and BAL (B) from ID#DHGF at day 7. (C) Quantification of total B cells from the blood and BAL over time. Dunnett’s multiple comparison test comparing to day 0. (D) Quantification of different B cell subsets in the blood and BAL as a frequency of total B cells. Showing the mean value from all animals with standard error. Dunnett’s multiple comparison test comparing values to day 0. No BAL data on B cells was collected for DGCX and DG3V. (E) Quantification of the frequency of IgG^+^, IgM^+^, IgA^+^, and isotype undefined of activated memory B cells. Dunnett’s multiple comparison test comparing values to day 0. DGCX and DG3V did not have baseline values. (F-H) Frequency of total B cells (F), subsets (G), and isotypes (H) from spleen, peripheral lymph nodes (axillary, inguinal, and/or cervical), previously hot pulmonary lymph nodes, normal pulmonary lymph nodes, and lung sections at day 10 post infection. No data from the lungs of DGCX, DG3V, DGRX, and DG4i; previously hot pulmonary lymph nodes from DGCX and DG3V; normal pulmonary lymph nodes from DGRX; or peripheral lymph nodes from DGRX. Significance for G and H done using a Turkey’s multiple comparison test. In G, significant difference in lung activated memory subset vs. spleen, peripheral LN, previously hot and normal pulmonary LN. Significant difference in activated naïve in lung vs. previously hot pulmonary LN. Significant difference in resting naïve in lung vs. spleen, peripheral LN, and previously hot pulmonary LN. In H, significant difference in IgG^+^ and unknown isotype in peripheral LN vs. spleen. Significant difference in unknown isotype in normal pulmonary LN vs. spleen. IgA isotype not quantified at day 10 necropsy. (I-K) Anti-spike IgG and IgM antibody responses in the plasma and BAL at day 0 and day 10 post infection. (I) Anti-Spike IgG antibody titration curves at day 0 and day 10 post-infection compared to uninfected and convalescent NHP plasma. Area under the curve (AUC) of the antibody titration curves for anti-Spike IgG and IgM responses at day 0 and day 10 in the plasma (J) and BAL (K). Significance calculated with Sidak’s multiple comparison test.

**Supplemental Figure 4 (refers to Figure 4 and 5).**
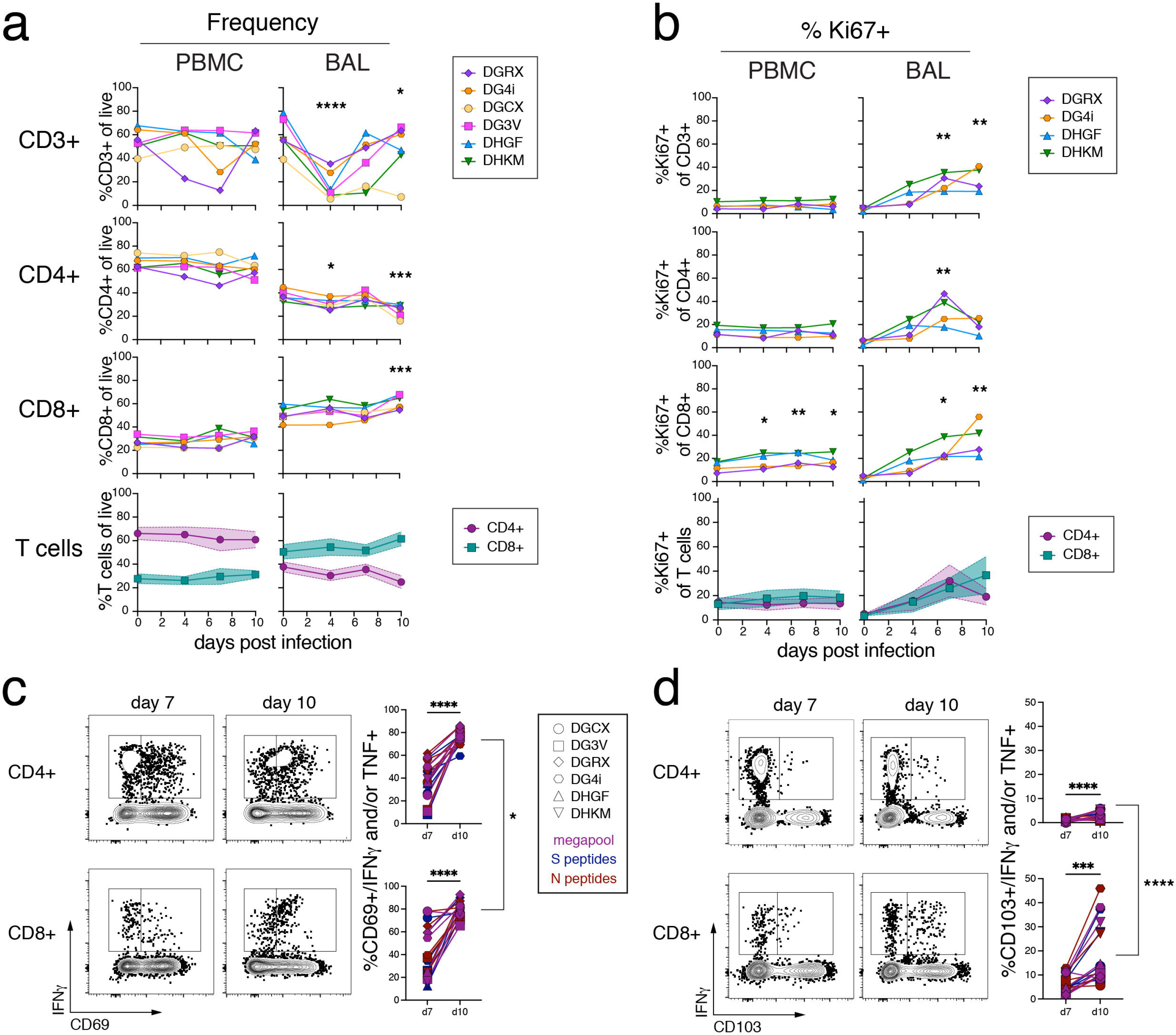
Activation of bulk and antigen-specific T cell responses in the blood and BAL. (A) Bulk CD3^+^, CD4^+^CD95^+^, or CD8^+^CD95^+^ responses overtime as a frequency of live cells in the blood and BAL. Significance indicated with Dunnett’s multiple comparison test comparing individual timepoint to baseline. Bottom set of graphs overlay the mean CD4^+^CD95^+^ and CD8^+^CD95^+^ response with standard deviation. (B) Frequency of Ki67^+^ on bulk CD3^+^, CD4^+^CD95^+^, or CD8^+^CD95^+^ in the blood and BAL. Bottom set of graphs overlay the mean Ki67^+^ on CD4^+^CD95^+^ and CD8^+^CD95^+^ response with standard deviation. Ki67 stain not done in DGCX or DG3V. (C) Representative flow cytometry plots from ID#DG4i and quantification of the frequency of CD69^+^ on Ag-specific CD4 and CD8 T cells from the BAL on day 7 and day 10 post infection. Paired t-test comparing day 7 vs. day 10 for CD4 and CD8 separately, and CD4 day 10 vs. CD8 day 10. (D) Representative flow cytometry plots from ID#DG4i and quantification of the frequency of CD103^+^ on Ag-specific CD4 and CD8 T cells from the BAL on day 7 and day 10 post infection. Paired t-test comparing day 7 vs. day 10 for CD4 and CD8 separately, and CD4 day 10 vs. CD8 day 10

**Supplemental Figure 5 (refers to figure 5).**
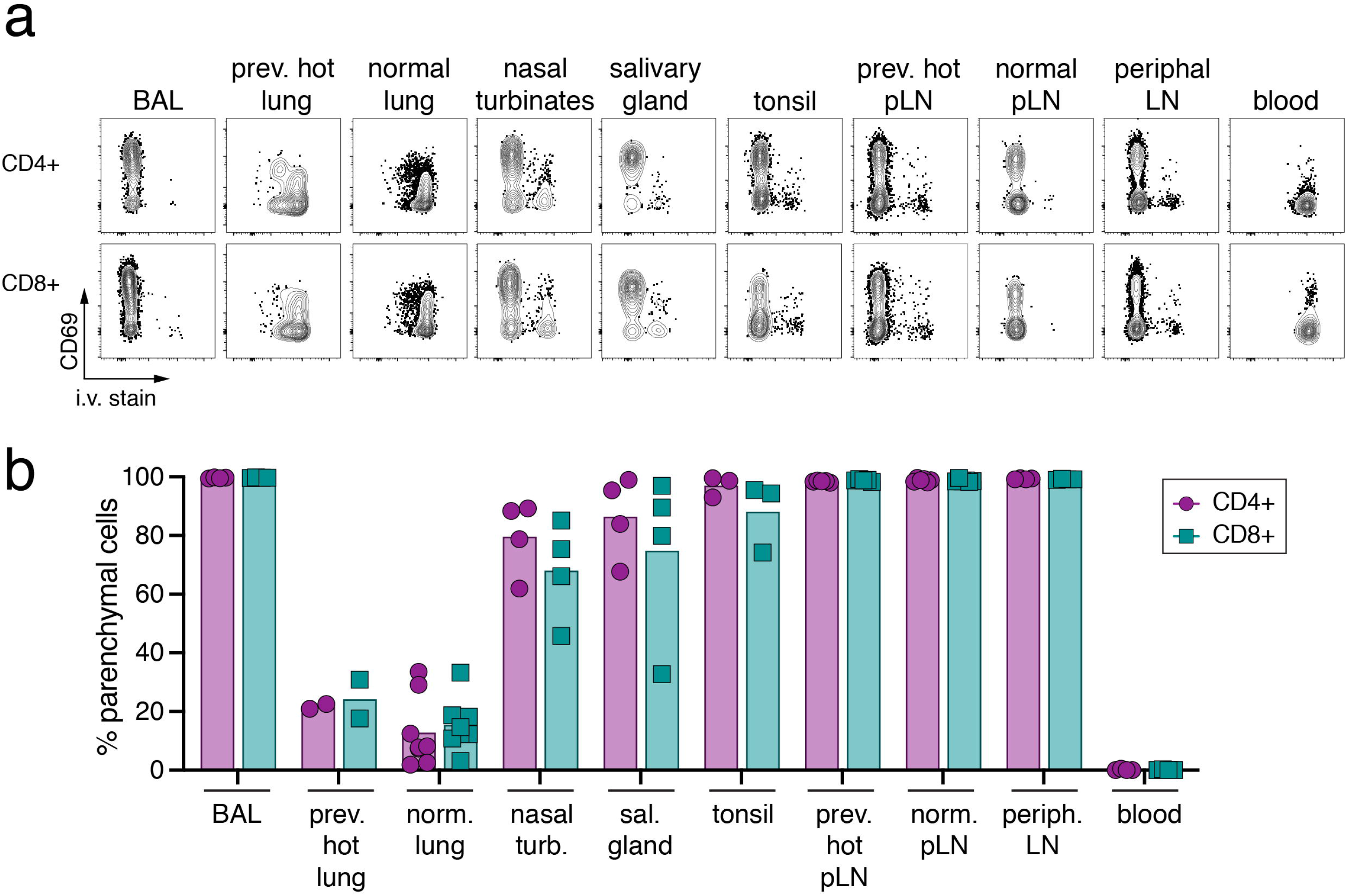
Parenchymal localization of T cells in tissues from SARS-CoV-2 rhesus macaques. (A) Representative flow cytometry plots of intravenous (i.v.) staining vs. CD69 on CD4^+^CD95^+^ and CD8^+^CD95^+^ from tissues at necropsy from ID#DGRX, except salivary gland, which was from ID#DGCX. (B) Quantification of parenchymal CD4^+^CD95^+^ and CD8^+^CD95^+^ T cells (equivalent to i.v. stain negative) from tissues at necropsy. DHGF and DHKM did not have i.v. stain.

